# Cross-feeding modulates the rate and mechanism of antibiotic resistance evolution in a model microbial community of *Escherichia coli* and *Salmonella enterica*

**DOI:** 10.1101/722561

**Authors:** Elizabeth M. Adamowicz, Michaela A. Muza, Jeremey M. Chacón, William R. Harcombe

## Abstract

With antibiotic resistance rates on the rise, it is critical to understand how microbial species interactions influence the evolution of resistance. We have previously shown that in obligate mutualisms the survival of any one species (regardless of its intrinsic resistance) is contingent on the resistance of its cross-feeding partners, setting the community antibiotic tolerance at that of the ‘weakest link’ species. In this study, we extended that hypothesis to test whether obligate cross-feeding would limit the extent and mechanisms of antibiotic resistance evolution. In both rifampicin and ampicillin treatments, we observed that resistance evolved more slowly in obligate co-cultures of *E. coli* and *S. enterica* than in monocultures. While we observed similar mechanisms of resistance arising under rifampicin selection, under ampicillin selection different resistance mechanisms arose in co-cultures and monocultures. In particular, mutations in an essential cell division protein, *ftsI*, arose in *S. enterica* only in co-culture. A simple mathematical model demonstrated that reliance on a partner is sufficient to slow the rate of adaptation, and can change the distribution of adaptive mutations that are acquired. Our results demonstrate that cooperative metabolic interactions can be an important modulator of resistance evolution in microbial communities.

**Significance statement:** Little is known about how ecological interactions between bacteria influence the evolution of antibiotic resistance. We tested the impact of metabolic interactions on resistance evolution in an engineered two-species bacterial community. Through experimental and modeling work, we found that obligate metabolic interdependency slows the rate of resistance acquisition, and can change the type and magnitude of resistance mutations that evolve. This work suggests that resistance evolution may be slowed by targeting both a pathogen and its metabolic partners with antibiotics. Additionally, we showed that community context can generate novel trajectories through which antibiotic resistance evolves.

## Introduction

The ability of pathogens to rapidly evolve antibiotic resistance is a pressing global challenge. While resistance frequently evolves in complex microbial communities, relatively little is known about how species interactions influence the evolution of antibiotic resistance (1–3). Most of the studies that do incorporate multiple species focus on the role of horizontal transfer of antibiotic resistance between species, and less on the *de novo* evolution of resistance within genomes (3–7). Additionally, antibiotic resistance studies in multi-species systems typically involve unknown interactions between species, with some exceptions in modelling (8–11). The specific role of interspecies interactions in the evolution of antibiotic resistance in microbial communities, therefore, remains largely unexplored.

Positive interactions are common in bacterial communities (12), and relatively understudied in terms of their role in modulating the evolution of resistance. One such interaction is cross-feeding, wherein two species exchange essential metabolites (13). The resilience of metabolically interdependent microbial systems to environmental disturbances is a growing field of study, with research being conducted into how these systems resist invasion (14, 15) and respond to abiotic environmental changes (8, 16–18). Over short time-scales (e.g. within a single growth curve), we previously showed that obligate cross-feeding produces ‘weakest-link’ dynamics (18), wherein the least resistant member of an obligate cross-feeding community constrains the ability of more resistant community members to grow at high antibiotic concentrations. In this case, the resistance (genetic mechanisms conferring an ability to grow at higher antibiotic concentrations) did not change for each species, but the tolerance (a phenotypic trait describing ability to grow at high antibiotic concentrations) was limited by the dependence on the least resistant species. This idea has also been demonstrated by others through modelling approaches (8).

We hypothesize that the weakest link pattern described above should also hold over evolutionary timescales; that is, at any given point during the evolution of resistance in a metabolically interdependent community, one ‘weakest-link’ species should set the tolerance of the whole community. The obligate cross-feeding interactions would then require that each individual species develop resistance for the whole community to rise in tolerance. We therefore hypothesize that metabolically interdependent communities will be slower to adapt to rising antibiotic levels than their single-species counterparts.

We also hypothesize that weakest link dynamics should affect the mechanisms of resistance evolution. For example, the evolution of some shared resistance mechanism such as an antibiotic-degrading enzyme (3, 19, 20) or an induction of tolerance mechanisms in a partner species (21, 22) could be uniquely selected for in co-culture. Cross-feeding may also limit the types of resistance mutations available to mutualistic networks of bacteria; for example, cross-feeding could make some resistance mechanisms more costly. Evolution of resistance by altering cell wall permeability may be particularly maladaptive in cross-feeding communities, where exchanged nutrients will typically be at low concentrations (23–25). Finally, a reduction in the rate of adaptation may drive different mechanisms of resistance to evolve. Varying the rate at which antibiotics are increased has been shown to lead to different evolutionary trajectories (23–28). More rapid changes in antibiotic concentration tend to select for mutations with larger effects that are more costly (23–28). These big effect mutations can trap populations on sub-optimal fitness peaks (26). We therefore hypothesize that we will see different mechanisms of resistance evolve in co-culture vs. monoculture, though the exact nature of these differences is unclear.

We sought to investigate whether obligate cross-feeding altered the rate and mechanism of antibiotic resistance evolution. We used a previously engineered two-species system of *E. coli* and *S. enterica*, wherein *E. coli* consumes lactose and excretes acetate that *S. enterica* uses as a carbon and energy source, and *S. enterica* overproduces methionine used by the methionine-auxotrophic *E. coli*. We evolved six replicate populations of each species growing in monoculture (providing *E. coli* with lactose and methionine, and *S. enterica* with glucose) and in obligately cross-feeding co-culture (providing both species with lactose only, hereafter referred to as ‘co-culture’) along increasing antibiotic gradients of rifampicin or ampicillin. At each transfer, the MIC (minimum inhibitory concentration) was assessed. We also constructed a mathematical model of resistance and used it to assess the generality of out findings. Our results show that growth in an obligate co-culture slows the rate of adaptation, and sometimes leads to different mechanisms of resistance.

## Results

### Antibiotic resistance evolves more quickly in monoculture than in obligate co-culture

First, we tested the rate at which antibiotic resistance evolved in monocultures of *E. coli* and *S. enterica* as well as obligate co-cultures of the two species. We established six replicate cultures of each monoculture, and the co-culture. Each culture was distributed along an antibiotic gradient of either ampicillin or rifampicin. After 48 hours of growth, we transferred cells in a 1/200 dilution to fresh medium in the same antibiotic concentration, as well as double that concentration (Figure 1A). We transferred populations for 20 transfers (approximately 180 generations) in rifampicin, and 10 transfers (approximately 90 generations) in ampicillin. At each transfer, we measured total population density (by OD600) and calculated MIC_90_ based on the density in wells along each gradient. After the final transfer, three colonies of each species were isolated from each replicate population and their MIC_90_ values measured. Due to contamination, we removed one replicate of rifampicin-evolved *S. enterica* in monoculture and only used data out to transfer 10 for the ampicillin-evolved lines.

**Figure 1.**
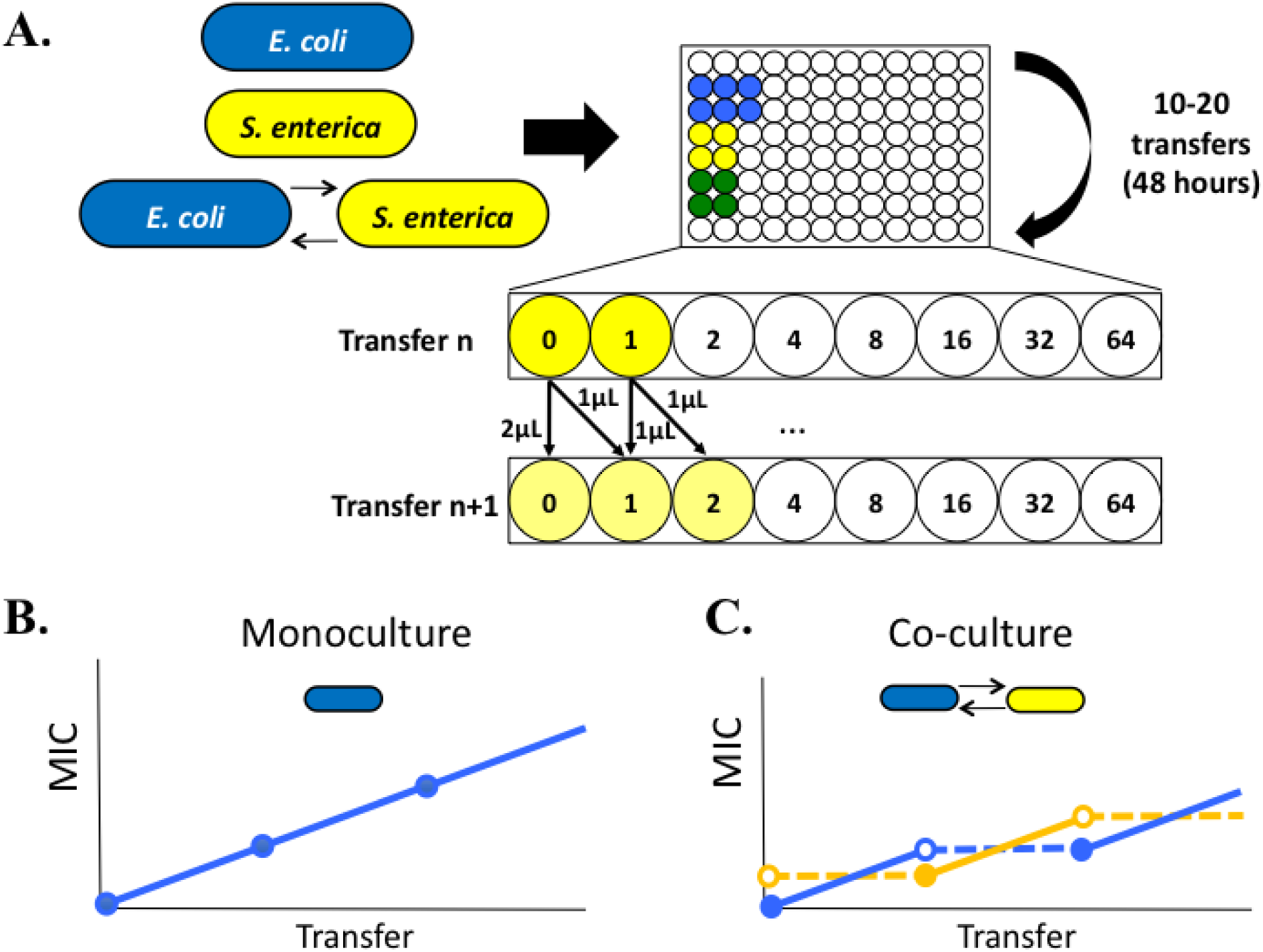
Schematic of experimental setup and expectations. **A.** Monocultures of *E. coli* and *S. enterica*, as well as cross-feeding co-cultures, were distributed in six replicate populations along an antibiotic gradient. The antibiotics tested included rifampicin and ampicillin, and the concentration of antibiotic increased twofold at each well. 96-well plates were incubated with shaking at 30°C for 48 hours, then cells were transferred to fresh medium and antibiotic in a new plate. The passaging regime was to transfer 1uL of culture to a fresh well containing the same concentration of antibiotic at which it had previously grown, and 1uL to a fresh well containing one concentration step higher antibiotic. At each passage, the OD600 of the plate was measured, as well as species-specific fluorescence (CFP for *E. coli*, YFP for *S. enterica*). **B-C.** Hypothesis on why time under selection may be sufficient to explain MIC differences between monocultures and co-cultures. **B.** In monoculture, each species is under selection at every time step, thus selecting for increasing resistance with each passage. **C**. In obligate co-culture, only the more antibiotic-sensitive species is under selection at a given time, and effective co-culture resistance requires an increase in MIC in both species. This leads to the slower rise in resistance for the co-culture.

We found that resistance evolved more quickly in monoculture populations than in co-cultures (Figure 2A,B). In rifampicin, growth in each monoculture was associated with a significantly higher increase in MIC per transfer than growth in co-culture (p = 0.0328 for *E. coli*, p = 0.0100 for *S. enterica*). In ampicillin, resistance in both species also rose more quickly in monoculture than co-culture, even over just ten transfers. The per-transfer increase in MIC of both monocultures was higher than that of co-cultures (p= 0.007 for *E. coli*, p= 0.0272 for *S. enterica*) (Figure 2B).

**Figure 2.**
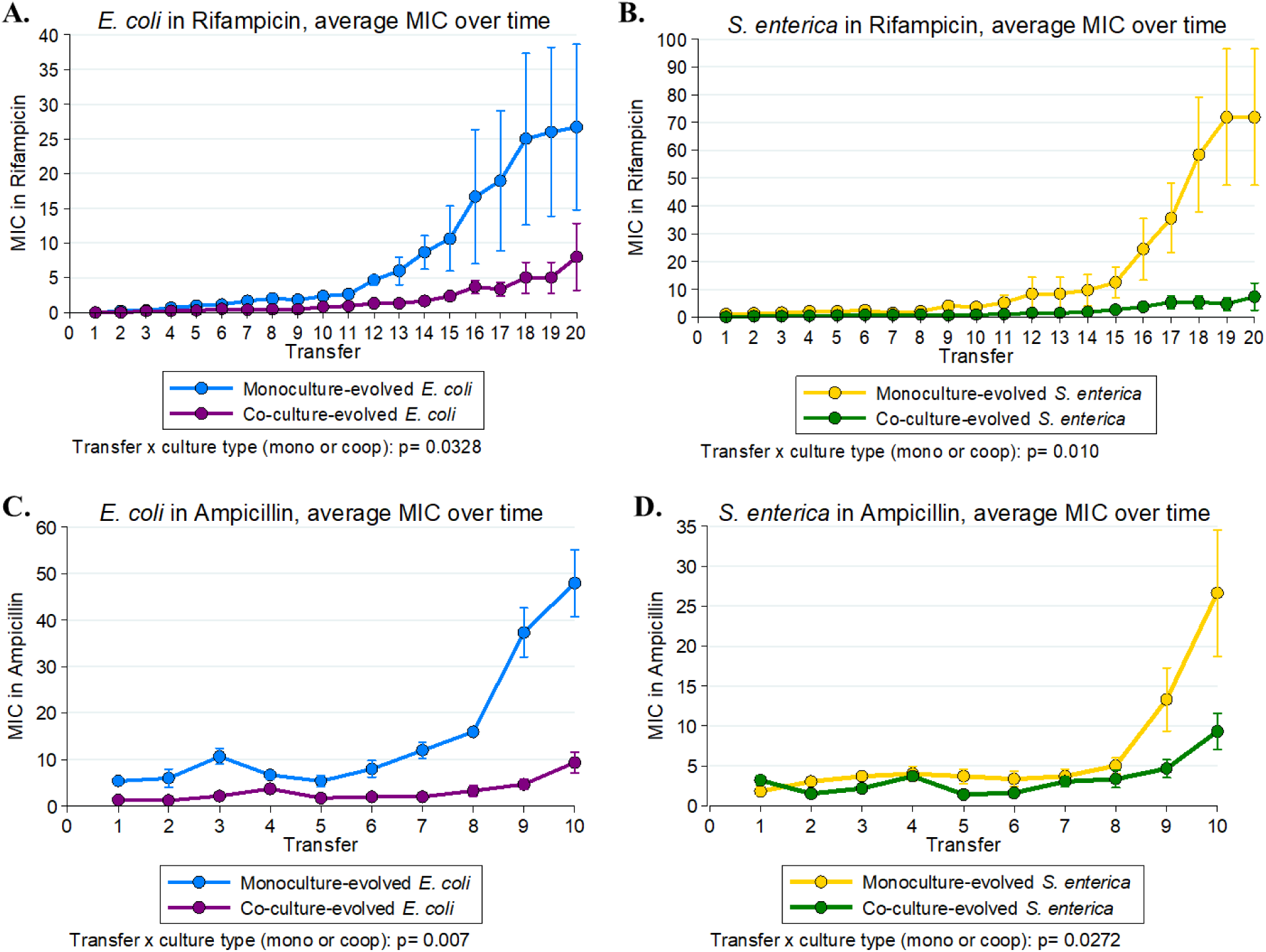
Resistance evolves more slowly in co-culture-evolved populations vs. monoculture-evolved populations. Six replicate populations each of monocultures and co-cultures were evolved along a rifampicin gradient (**A-B**) or an ampicillin gradient (**C-D**). Population MICs for each species (*E. coli* **A, C**; *S. enterica* **B, D**) were measured each transfer and the resulting MICs plotted. Statistical analysis was performed using a mixed-effects model with a randomized slope for each replicate within a culture type. P-values are for the interaction term between passage and culture type.

### Similar mechanisms of rifampicin resistance evolve in monoculture and obligate co-culture

To determine whether cross-feeding selected for different resistance mechanisms we sequenced resistant populations at the end of the experiments. Genomic DNA was extracted from the well that grew at the highest concentration of antibiotic for each replicate population. For each resistant population we created a list of mutations excluding any mutations also observed in antibiotic free controls (**Supplementary Tables 1 and 2**). For further analysis, we focused on genes that acquired mutations multiple times within a treatment, as parallel evolution is a signature of adaptation (27).

Under rifampicin pressure, the genes that acquired mutations in co-culture were a subset of those that acquired mutations in monoculture for both species (Figure 3A,B). The most clearly identifiable resistance-associated mutation was *rpoB*, a component of RNA polymerase and the most common mutational target for rifampicin resistance (28). Mutations in this gene arose in four out of the six replicates each in monoculture and co-culture (Figure 3A) and were strongly tied to higher levels of resistance (Figure 3C,D). A mutation in a prophage tail-specific protein, *prc*, was also repeatedly observed in both monoculture and co-culture (**Supplementary figure 1**). The overlap in mutations suggest that co-cultures and monocultures are evolving along similar adaptive trajectories.

**Figure 3.**
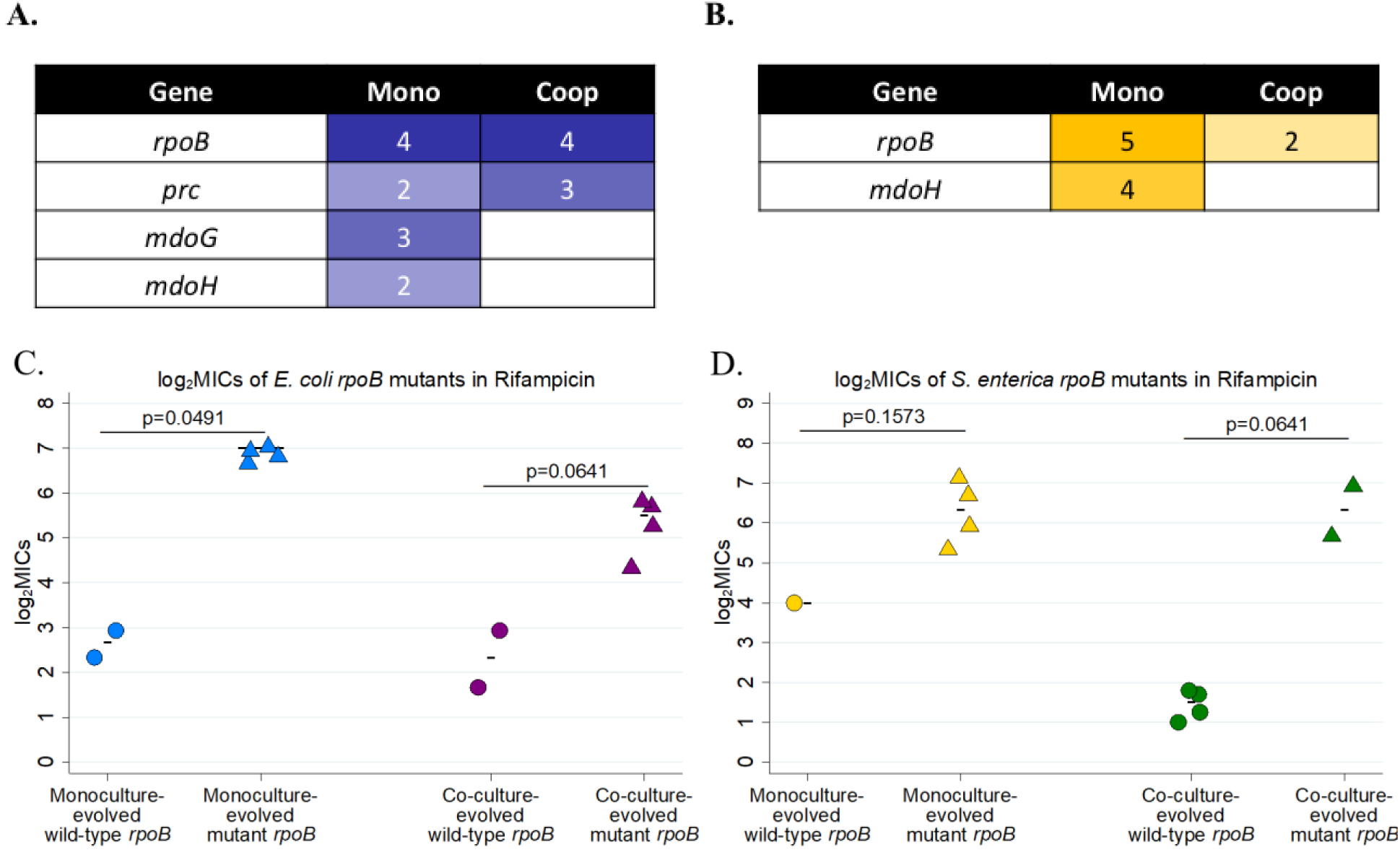
Resistance-associated mutations in rifampicin-resistant evolved populations **A-B.** Lists of mutations which arose in *E. coli* (**A**) or *S. enterica* (**B**) monocultures and co-cultures. The number in each box represents the number of independent replicates in which the putative resistance mutation was observed. Additional mutations that occurred in only one population may be found in **Supplementary table 1**. **C-D.** Rifampicin MICs for isolates with wild-type vs. mutant *rpoB* genes in *E. coli* (**C**) and *S. enterica* (**D**). Isolates were obtained from passage 20 populations by streaking onto selective medium and picking isolated colonies. MIC_90_ values for isolates were defined as the lowest concentration of antibiotic which decreased growth by greater than 90% by 48 hours at 30°C. Each point represents the average MIC of three isolates from a single population.

Monoculture lines evolved more mutations than co-culture lines. Mutations in *mdoG* and *mdoH* were only observed in monocultures of *E. coli* and *S. enterica* under rifampicin selection. Both *mdoG* and *mdoH* likely influence cell membrane permeability (29, 30). The mutations were not associated with any changes in monoculture or co-culture growth rates (**Supplementary figure 2**). Taken together, the pattern of rifampicin resistance mutations suggests that populations were moving along the same evolutionary trajectory in monoculture and co-culture, but got further along that trajectory in monocultures.

### Different mechanisms of ampicillin resistance arise in monocultures vs. obligate co-cultures

In contrast to our results from rifampicin, the mutational spectra that we obtained from sequencing ampicillin-resistant populations suggests different resistance mechanisms arose in monoculture vs. co-culture. We identified more mutations in ampicillin than in rifampicin due to our lowered threshold for inclusion (above 50% frequency, verses above 80% in rifampicin); we used this cut-off to be able to detect resistance mutations even with the fewer number of transfers in ampicillin. Nevertheless, we observed distinct mutational signatures in both *E. coli* and *S. enterica* monocultures and co-cultures.

Very few genes acquired mutations more than once in the *E. coli* replicates. Two co-cultures replicates evolved mutations in *proQ*, a regulator of efflux pumps (31) (Figure 4A). In monoculture evolution repeatedly involved a gene associated with stress response (*rne*). Given the relative paucity of mutations there is at best only a weak signature of differential resistance mechanisms evolving in *E. coli* in monoculture and co-culture.

**Figure 4.**
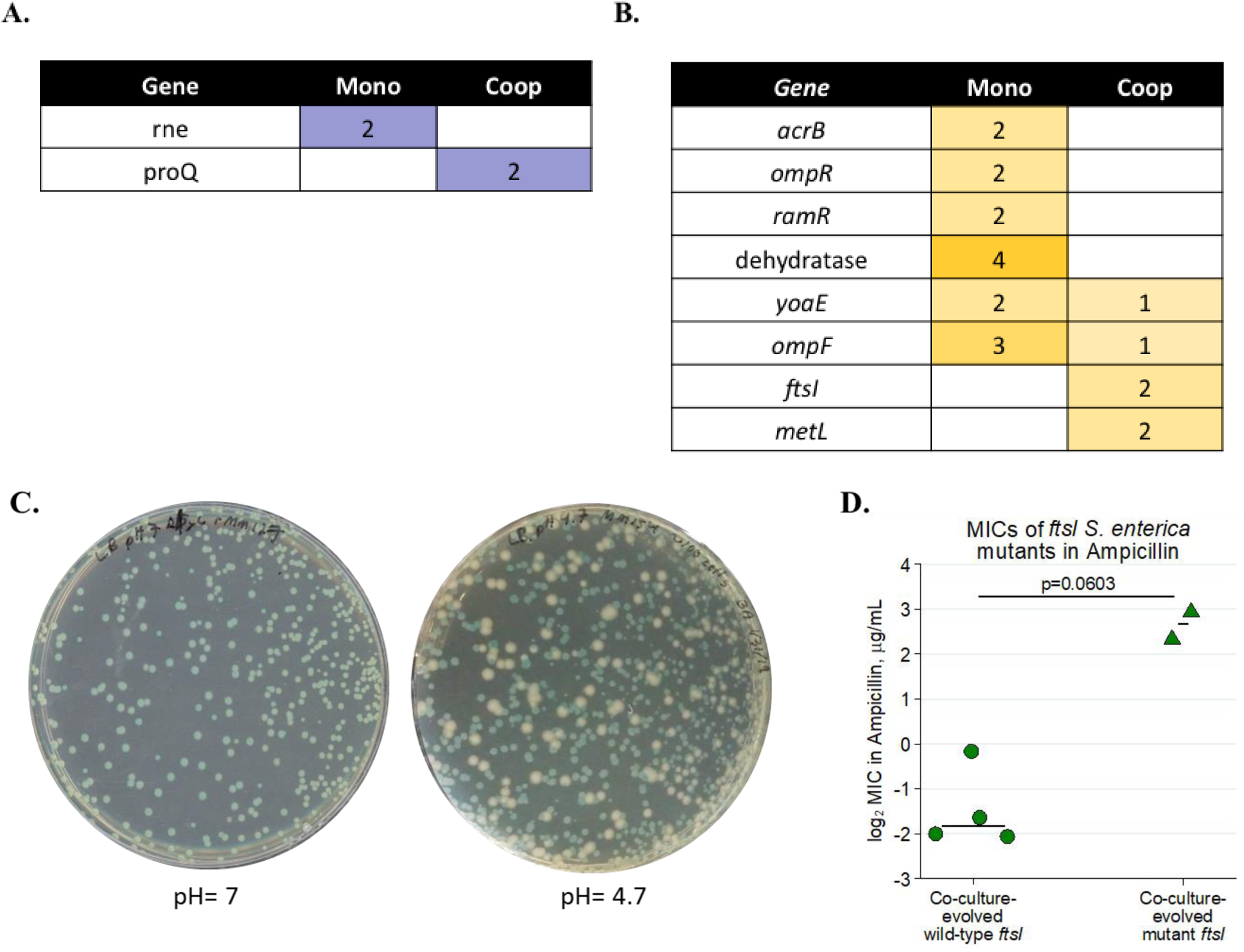
Resistance-associated mutations in ampicillin-resistant evolved populations **A-B.** Lists of mutations which arose in *E. coli* (**A**) or *S. enterica* (**B**) monocultures and co-cultures. The number in each box represents the number of independent replicates in which the putative resistance mutation was observed. Additional mutations that occurred in only one population may be found in **Supplementary table 1**. **C.** Image of co-culture populations containing *ftsI* mutations on Petri plates with LB pH=7 (left) and pH=4.7 (right). Blue colonies are *E. coli* which metabolize X-gal to a blue color; white colonies are *S. enterica*. No white colonies were observed on LB at pH=7. **D.** MICs of isolates from libraries containing *ftsI* mutations in pH=4.7 medium. Isolates were obtained from passage 10 populations by streaking onto selective medium and picking isolated colonies. MIC_90_ values for isolates were defined as the lowest concentration of antibiotic which decreased growth by greater than 90% by 48 hours at 30°C. Each point represents the average MIC of three isolates from a single population. P= 0.0603, Mann-Whitney U.

The evidence for differential mechanisms of resistance was stronger in *S. enterica*, despite more overlap in mutated genes (Figure 4B). Mutations in efflux pump genes (*acrB, ramR*), cell permeability (*ompR*) and an unnamed dehydratase, arose in *S. enterica* monoculture but not in co-culture. None of these mutations were associated with a significant increase in resistance except for *ramR*, wherein mutants had a higher median MIC than wild-type isolates (p=0.0491, **Supplementary figure 3**). Conversely, mutations in *ftsI*, a penicillin-binding protein, was only observed in co-culture. *FtsI* is thought to be an essential gene and is a penicillin-biding protein (PBP3), which is known to be a target site for β-lactam antibiotics (32). Interestingly, the mutations we observed in *ftsI* in our whole-genome sequencing were a combination of point mutations (D534Y at 75% frequency in rMM158) and mutations which should have ablated gene function (+A at 100% frequency in rMM127 and Q142* in rMM158). *S. enterica* isolates were difficult to obtain from these populations, and isolates which we did manage to isolate either did not contain the +A or Q142* mutations, or had a suppressor mutation in the same codon as the Q142* mutation site which eliminated the stop codon (**Supplementary table 1)**.

Castanheira et al. recently demonstrated that the lethality associated with *ftsI* loss in *S. enterica* could be mitigated by growing *S. enterica* under acidic conditions (pH<5.8); under these conditions, a second PBP3, which they called PBP3_SAL_, is expressed (33). In light of this, we plated co-culture populations containing Q142* or +A mutations in *ftsI* on LB of pH 4.7. On acidic plates we were able to obtain *S. enterica* isolates at roughly equal frequencies to what we would expect in the population, and Sanger sequencing demonstrated that these isolates did contain the loss-of-function Q142* or +A mutations. These isolates did not show detectable growth in monoculture unless the growth medium was acidified (Figure 4C), and were associated with increased MICs verses isolates with wild-type *ftsI*, though this difference was not statistically significant (Figure 4D, p= 0.0603). Interestingly, *ftsI* mutant isolates had co-culture growth rates comparable to wild-type when paired with an *E. coli* ancestor (**Supplementary figure 4**). This suggests that *ftsI* knockout mutations were non-viable in monoculture but conferred little cost in co-culture.

### A model suggests that interdependency is sufficient to generate differences in evolution of resistance

To test the effect of interdependence on evolution of antibiotic resistance in the absence of species-specific biological details, we developed a simple model. The model is based upon the haploid Wright-Fisher model with selection. Populations were kept at a fixed density with a transfer regime similar to our experiments. Individuals had a gene governing MIC which could mutate upon reproduction and determined whether the individual could grow in a well with a given antibiotic concentration. In co-culture simulations species were interdependent, such that no individuals of a species could grow if there were no viable individuals of the other species in a well (see Methods for details).

We first examined the relationship between the number of interdependent species and the rate of evolution of antibiotic resistance. MIC-increasing mutations were pulled from a uniform distribution ranging from zero to 1.1 (with 1 being the step-size at which antibiotic concentration increased across wells). One thousand individuals of each species were simulated per well. Ten generations occurred within a transfer, and we conducted nine transfers. Thirty-five replicates were simulated. We found that antibiotic tolerance (the highest antibiotic concentration at which a species grew) evolved more slowly as more interdependent species were simulated (Figure 5A, interaction between # species and transfer in linear mixed-effects model, t = −9.4, p < 1e-7). When the mutation rate or the population size was smaller, cross-feeding more strongly reduced evolution of resistance (Figure 5B, one-way ANOVA, F(2,297) = 6.9, p = 0.0012). More mutations accumulated in resistance populations when there were fewer species present (**Supplementary figure 5**, F(2,102) = 20.98, p < 1e-7).

**Figure 5.**
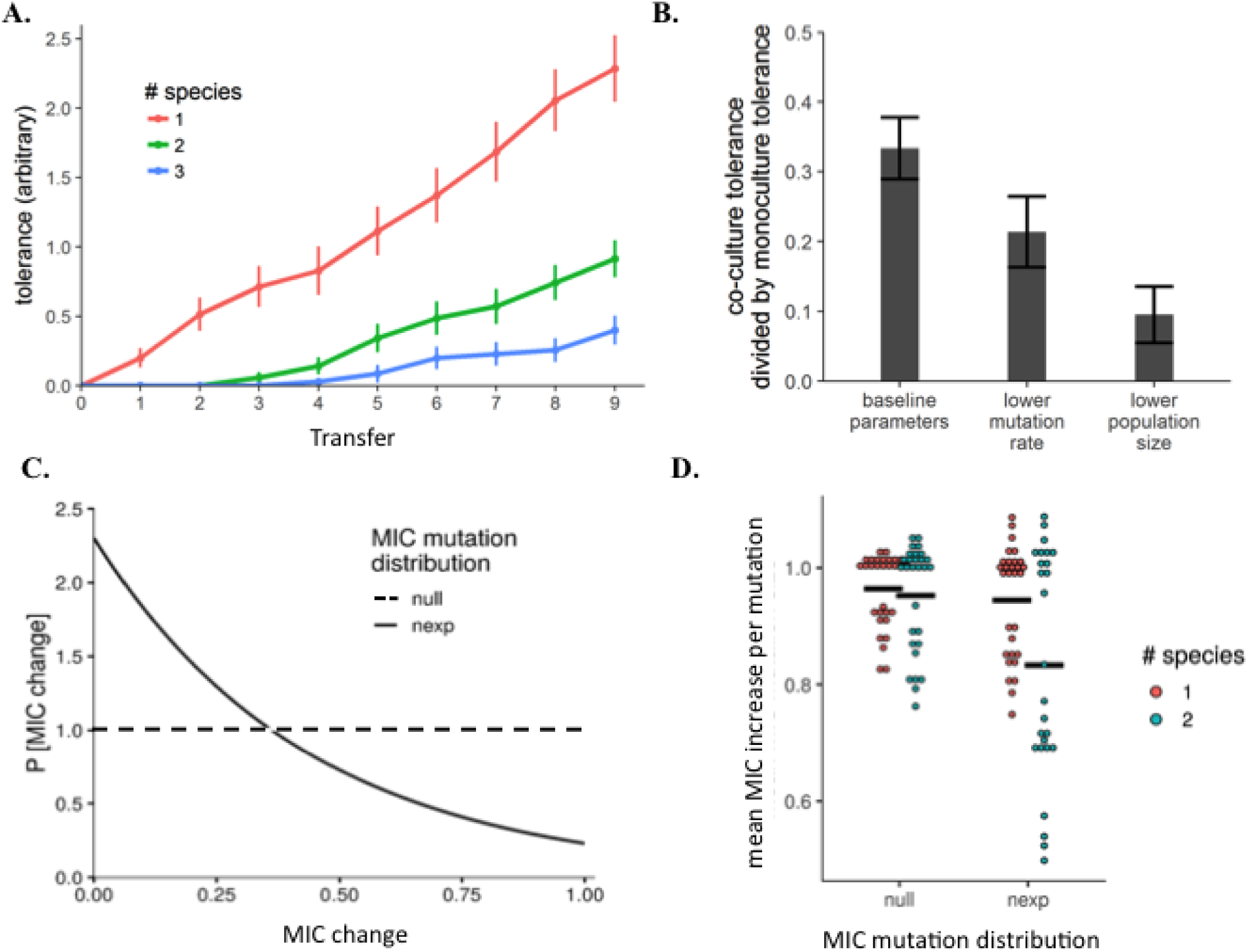
Simulation model of the evolution of antibiotic resistance in single-species vs. multi-species obligately dependent communities. **A.** A simulated evolution experiment with 35 replicate populations shows that increasing the number of interdependent species in a consortium results in a slower increase in the average MIC over time. **B.** Effect of lower mutation rate and lower population size on resistance evolution in model. Both lower mutation rates and lower population sizes result in bigger differences in the relative tolerance of one-species versus two species systems. **C.** Distributions from which MIC-altering mutations were randomly pulled. The x-axis is relative to the maximum MIC change possible in one mutation. The “null” model used a uniform distribution. The “nexp” used a truncated negative exponential distribution with rate parameter 2.3, which kept 90% of random numbers less than the max MIC change of 1.0. **D.** Mean MIC increase conferred per mutation as a function of the mutation distribution and the number of species.

We next tested how the distribution of mutations influenced the mechanisms of resistance. We simulated evolution when all mutations were equally likely versus when mutations which conferred a greater resistance were less common (Figure 5C). Under a uniform distribution there was no difference in the average size of resistance mutations accumulated in monocultures and co-cultures (Figure 5D). However, when big-effect mutations were rare populations in co-cultures acquired significantly smaller effect mutations than populations in monoculture (Figure 5D, two-way ANOVA, interaction between # of species and mutant distribution: F(1,111) = 5.087, p = 0.026; Tukey’s HSD finds only nexp + 2 species is different from other treatments). Our results demonstrate that when the frequency of mutations is not uniform, populations in monoculture and co-culture will sample different sets of mutations.

## Discussion

Our results demonstrate that obligate cross-feeding changes the rate and mechanisms of resistance evolution relative to monoculture. We showed that the obligate interspecies interactions resulted in a slower rate of adaptation to two different drugs and in an evolution model unbounded by species-specific details. In rifampicin, the resistance mechanisms remained similar between monoculture and co-culture-evolved lines; genes that acquired resistance mutations in co-culture were a subset of the genes involved in adaptation in monoculture. In ampicillin, in contrast, we observed different resistance mechanisms arising in monoculture and in co-culture, likely due in part to different costs of resistance mutations in monoculture and co-culture. In particular, we showed that *ftsI* mutations provide high levels of ampicillin resistance, but are only selected for in co-culture. Overall, our experimental and modeling results showed that monocultures evolved more resistance than co-cultures, irrespective of species and antibiotic identity, suggesting that slower resistance evolution in cross-feeding co-cultures is a general phenomenon.

We show that mutualistic interactions between species can result in slower evolution of antibiotic resistance. This demonstrates the importance of taking ecological context into account when studying antibiotic resistance, and suggests that species interactions may play an important role in shaping resistance evolution trajectories. It is important to note that populations in co-culture experienced different physiology and population sizes, which could have influenced rates of adaptation (34). However, our mathematical model demonstrates that interdependency is sufficient to drive differences in rates of adaptation to abiotic stressors, independent of other biological factors. Our results suggest that using a broad-spectrum antibiotic that targets both a pathogen and its obligate cross-feeding partners would slow the rate at which pathogens acquire resistance.

We found many resistance mechanisms that evolved repeatedly. As expected, rifampicin treatment led to evolution of *rpoB* mutations in both monocultures and co-cultures (26, 28, 35–37). We also routinely observed mutations in genes associated with cell wall permeability. Genes that influence cell wall biochemistry (*mdoG* and *mdoH*) repeatedly evolved in rifampicin, while porin and efflux pump genes (*ompR* and *ompF*) evolved in ampicillin. Modulation of cell wall permeability through changes in porin regulation (38, 39) and expression of efflux pumps (40) are well-established resistance mechanisms. However, less is known about how modification of cell membrane biochemistry affects resistance. Changes in membrane phospholipid composition was recently shown to increase *Staphylococcus aureus* resistance to daptomycin (41), and changes in LPS biosynthesis in *E. coli* increased sensitivity to bacitracin and rifampicin (30). Our work adds to the evidence that resistance can repeatedly evolve through diverse mechanisms.

The *ftsI* mutation we identified in *S. enterica* only arose in co-culture environments, despite being associated with increased ampicillin resistance. This is particularly interesting as *ftsI* activity is implicated in interactions between *S. enterica* and the human immune system (33). Loss of *ftsI* might provide high-level resistance but prevent *S. enterica* from reproducing outside the acidic macrophage environment, making *S. enterica* into an obligate intracellular pathogen. Antibiotic pressure in a patient might therefore select for antibiotic-resistant *ftsI* mutants which are unable to spread. Other studies have demonstrated that resistance mutations have associated fitness costs in specific environments (42, 43). However, further research is needed into how these context-dependent costly mutations might be disproportionately selected for in multispecies environments where those costs are not realized until an infection situation.

Our mathematical model suggests an additional reason that different mechanisms of resistance are likely to be observed in monoculture and co-culture. Experimentally, we observed that growth in co-culture changed the cost of a resistance mutation. Our model suggests that the benefit of resistance mutations is also likely to change in different growth contexts. In agreement with previous work (35, 44) evolution in monoculture simulations was driven by the mutations that caused the biggest MIC impact, even if they were rare. In contrast, in co-culture, resistance evolution occurred primarily through mutations of smaller effect. This outcome makes sense as there is little benefit to evolving resistance beyond that of obligate partners. If small effect mutations are more common and provide the same benefit as big effect mutations, then small effect mutations are more likely to be observed in evolutionary trajectories. Future studies testing whether this effect is observed when experimental sample sizes are increased will shed further light on this hypothesis.

Studying how mutualisms evolve resistance to environmental stresses has significant implications not only for the evolution of antibiotic resistance in microbial communities, but also for the impact of climate change on symbiotic relationships at the macroscopic level. Our results suggest that organisms involved in obligate mutualisms, such as plant-pollinator interactions, may be constrained in their ability to adapt to a changing climate. Microbial communities offer an intermediate between theoretical models and long-term macroscopic mutualism studies. This allows us to test model predictions in a more reasonable time frame and potentially develop a predictive framework for the coevolution of mutualisms that extends to macroscopic communities. Overall, this work emphasizes the need to take ecological context (and particularly partner resistance) into account when studying the rates and mechanisms through which populations adapt to abiotic challenges.

## Methods

### Bacterial Strains and Media

The *Escherichia coli* and *Salmonella enterica* strains used in this study have been described previously (45). Briefly, the *E. coli* str. K12 was the Δ*metB* strain from the Keio collection (46) that was mated with an hfr line to reinsert the lac operon (29). The *S. enterica* LT2 mutant was selected and engineered to excrete methionine (47). Each strain is fluorescently labelled with a genomic integration of a fluorophore; *E. coli* is labelled with cyan fluorescent protein (CFP) and *S. enterica* is labelled with yellow fluorescent protein (YFP). Bacteria were grown in minimal Hypho medium containing phosphate and varying amounts and types of carbon and nitrogen as previously described (45). Co-culture media contain 2.78mM of lactose, *E. coli* monoculture medium 2.78mM of lactose and 0.020mM of methionine, and *S. enterica* medium contained 5.55mM of glucose.

### Experimental Evolution

Three different culture conditions (*E. coli* monoculture, *S. enterica* monoculture, and obligate co-culture) were each evolved across two antibiotic gradients (rifampicin, Chem-Impex International Inc. 00260, and ampicillin, Fisher BP1760). Six replicate populations were evolved for each antibiotic-culture condition combination. Each antibiotic gradient began with an antibiotic-free control well and increased twofold with each subsequent well, starting at 0.25 µg/mL and ending at 64 µg/mL. When populations evolved to grow at 64µg/mL, the upper end of the gradient was increased to 1024µg/mL and the lowest concentrations were removed. To minimize differences in concentration between transfers caused by pipetting errors, stocks of antibiotic gradients were prepared in advance and diluted such that 2µL of stock constituted the desired concentration when diluted 1/100 in the bacterial growth plate. Fresh antibiotic stocks were prepared and filter-sterilized immediately before the experiment began.

Initial bacterial cultures were inoculated from DMSO freezer stocks in 10mL of monoculture minimal Hypho medium for approximately 48 hours at 30°C, to stationary phase (OD~0.4). Cells were then distributed into 96 well cell culture plates. For monocultures, 2µl of bacterial cells (~2×10^5^ cells) were inoculated into 196µl of the appropriate monoculture medium with 2µL of antibiotic stock. In co-cultures plates, 1µl of *E. coli* and 1µl of *S. enterica* were inoculated into 196µL medium. The plates were then incubated for 48 hours at 30°C with shaking at 450 rpm. After each growth phase, cells were then transferred to a new 96-well plate with fresh media. 1µL cells were transferred to a new well with the same antibiotic concentration, and 1µL cells were transferred to a new well that was one step higher on the antibiotic gradient (see Figure 1). This regime challenged bacterial populations with increasing antibiotic concentrations at each transfer, but also allowed for populations to be maintained throughout the experiment if resistance was not acquired during a given transfer. The 96-well plate from each growth phase was frozen down in 10% DMSO for future analysis. After each 48-hour growth period, each plate was also placed onto a Tecan InfinitePro 200 plate reader where OD600 and species-specific fluorescence measurements were obtained. These readings were used to calculate the 90% minimum inhibitory concentration (MIC_90_) for each replicate; growth at a given antibiotic concentration was confirmed if the well OD600 was above 10% of the OD600 of the antibiotic-free control well for a given replicate. Statistical analysis of rate of resistance evolution between monocultures and obligate co-cultures for each antibiotic was performed in R. A linear mixed-effects model with a random slope for each replicate within a treatment group was used to test the relationship between MIC and transfer, culture type (monoculture or co-culture), and a transfer-culture type interaction.

### Sequencing

To identify mutations which conferred antibiotic resistance in each evolved population, the most resistant population of each replicate in each antibiotic-growth condition combination was whole-genome sequenced. Two antibiotic-free populations per antibiotic-growth condition combination were also sequenced to identify any mutations which may have arisen during passaging but are not related to antibiotic resistance. Each population to be sequenced was scraped from 10% DMSO freezer stock and grown up in 10mL Hypho at the appropriate antibiotic concentration for 48 hours at 30°C. gDNA was then extracted from each population using Zymo Quick-gDNA Miniprep Kit (11-317C). The gDNA was then used to prepare Illumina sequencing libraries according to the Nextera XT DNA Library Prep Kit protocol. Libraries were submitted to the University of Minnesota Genomics Center for QC analysis and sequenced on an Illumina Hi-Seq with 125bp paired-end reads.

Sequence analysis was performed using BreSeq (48) to align Illumina reads to reference *E. coli* and *S. enterica* genomes as previously described (49). Briefly, mutation lists for resistant populations were filtered such that variation between our ancestral strains and the reference genome were removed, as well as any mutations which also arose in the antibiotic-free populations. Mutation lists were then assembled for each population (**Supplementary table 1**) and any known functions described (**Supplementary table 2**). Mutations which rose to above 50% in frequency were examined further for their possible role in conferring antibiotic resistance (**Table 1**).

### Isolation and Phenotyping of Isolates

#### Obtaining isolates

At the final transfer of each evolution experiment, the well in each replicate displaying growth at the highest antibiotic concentration, according the MIC_90_, was identified as the ‘resistant population’. These resistant populations, as well as the corresponding antibiotic-free control populations from each replicate, were plated onto minimal Hypho medium containing X-Gal (5-Bromo-4-chloro-3-indolyl β-D-galactopyranoside, Teknova X1220), which allowed us to differentiate between β-galactosidase-positive *E. coli* and β-galactosidase-negative *S. enterica* through a colorimetric change. Three isolates of each species each from the resistant and antibiotic-free wells were selected from these plates and frozen down in 10% DMSO for further analysis.

For ampicillin-evolved populations in which we were unable to obtain *S. enterica* isolates on pH neutral medium, we prepared acidic Hypho plates by adding 6M HCl to the growth medium until the pH reached ~4.7. Growth medium was then autoclaved and X-gal was added as described above. For growth of these isolates in liquid medium, 6M HCl was added to autoclaved Hypho medium to a pH of ~4.7.

#### Isolate growth rates and yields

Evolved isolates were scraped from DMSO freezer stocks onto solid agar plates and grown up overnight, then inoculated into 96-well plates containing the appropriate species-specific Hypho and grown at 30°C for 48 hours at 450rpm to acclimatize them to minimal media growth and standardize cell density. 2µL of stationary-phase cells was then used to inoculate a new 96-well plate containing 198µL monoculture growth medium. These plates were then placed in a Tecan InfinitePro 200 for 48 hours at 30°C with shaking at 300rpm; OD600, CFP, and YFP were measured every 20 minutes. After 48 hours, 1µL from the wells of monoculture isolate plates were transferred to new 96-well plates. These plates contained 198ul of fresh co-culture Hypho medium and 1ul of either ancestor strain of *E. coli* or *S. enterica.* If the monoculture isolate was *E. coli*, the ancestor strain of *S. enterica* was added, and vice-versa. The plates were then placed into a Tecan InfinitePro 200 for 48 hours at 30°C, and OD600 and florescent data was obtained every 20 minutes. Growth rates and yields were then calculated using Baranyi curves in R software using an in-house script. Statistical analyses and graphs were prepared in Stata version 14.

#### Isolate Minimum Inhibitory Concentrations (MICs)

Evolved isolates were prepared for inoculation as described above for beginning monoculture growth rate experiments. Based on the MIC_90_ of the population from which they originated, 2uL of each isolate culture was then inoculated across an antibiotic gradient such that the antibiotic concentration equaling the population MIC_90_ was in the middle of the gradient. An antibiotic-free control well was also included for each gradient. If the gradient used was insufficient to calculate MIC (e.g. cells grew at all concentrations, or at none of them), the experiment was repeated with the antibiotic gradient shifted up or down as necessary. MIC of isolates was determined using MIC_90_ as described above. Statistical analyses and graphs were prepared in Stata version 14.

### Evolution Model

We used an evolution simulation to examine the rate at which antibiotic resistance evolved as a function of the number of interdependent species and other variables. There were two time scales in this model. The “within-transfer” time scale, and the “between-transfer” time scale. Within a transfer, evolution was simulated in any given “well” using a modified haploid Wright-Fisher simulation with selection. A well was initiated with N individuals (default N = 1000). If this was the first transfer, these individuals were clones of a genotype with growth rate (s) = 1 and MIC = 0. The well had a pre-determined antibiotic concentration. Evolution occurred over a predetermined number of generations. Each generation, the population was fully replaced. The new population was picked from the previous generation depending on the frequency of each genotype and its growth rate. The unscaled fitness of individual *i* was determined by 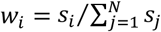. This was scaled to get expected frequency 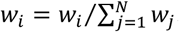. In other words,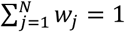 The next generation was then created by generating N random numbers from a uniform distribution on [0,1] and choosing genotypes from the previous generation based on their *w*_*j*_. Prior to calculating *w*, each genotype’s MIC was compared to the antibiotic concentration in the well. If the genotype’s MIC was less than the antibiotic concentration, its *s* = 0 for the fitness calculation.

Once the new population was generated, some of the new individuals may be mutated. Each simulation used a pre-determined mutation rate *u* (default = 0.001 mutations per individual per generation). We generated N random numbers from a uniform distribution on [0,1]. Any random numbers less than *u* meant those individuals gained a mutation. In all cases, mutations could be either a growth rate mutation or an MIC mutation with a 50:50 chance. Growth rate mutations were simulated by adding a random number from a normal distribution with mean 0 and standard deviation 0.1 to the current growth rate (i.e. deleterious growth rate mutations were equally as likely as advantageous). MIC mutations altered the MIC of the individual, as described in the results. We tracked the genealogy of all individuals.

To simulate interdependence, we simulated >1 species per well. Each species had N individuals, and the dynamics of each species were independent of the other species, *except* that there must be individuals of all species with an MIC > the antibiotic concentration for any species to grow.

The second time scale—the transfer scale—was simulated to approximate the wet-lab experiments. Individuals were transferred from one well to two wells: the same antibiotic concentration, and the next higher antibiotic concentration. Therefore, wells received individuals from two wells. If there were surviving species in both source wells, N/2 individuals were randomly chosen from each well to populate the new well. If there were surviving species in only one source well, all N individuals were transferred from that well to the new well. If no source wells had surviving species, the destination well was sterile. Simulations were conducted in R.

Statistical analyses of the simulation data: We analyzed the rate of evolution of antibiotic tolerance as was done in the wet-lab experiments. A linear mixed-effects model with a random slope for each replicate within a treatment group was used to test the relationship between the response variable MIC and explanatory variables transfer, culture type (monoculture or co-culture), and a transfer-culture type interaction. A one-way analysis of variance tested the amount of mutations observed in the most-tolerant population, with an independent variable of the # of interdependent species. To calculate the relative tolerance of two-species simulations versus one-species simulations (Fig. 5C), we first found the mean and standard error in the tolerance in each set of simulations. Then, the mean of the two-species community was divided by the mean of the one-species community. The error was propagated using (mean_relative_tolerance)*sqrt((SEM[two species]/mean_tolerance[two_species])^2 + (SEM[one species]/mean_tolerance[one_species])^2). The average MIC mutation size was the (Fig. 5D) was average MIC in a surviving population divided by the average number of mutations in that population. In all cases, since species were functionally equivalent, we arbitrarily chose one species to assess. Statistics were performed in R. Example code to run simulations is available as supplementary files.

## Author contributions

E.M.A, M.A.M., J.M.C. and W.R.H. designed research; E.M.A, M.A.M., and J.M.C. performed research; E.M.A., J.M.C., and W.R.H. analyzed data; and E.M.A., J.M.C., and W.R.H. wrote the paper.

## Acknowledgements

The authors would like to thank Lisa Fazzino, Sarah Hammarlund, Brian Smith, and Leno Bernard Smith Jr. for their insights on this work. This work was supported by a Natural Sciences and Engineering Research Council of Canada Postgraduate Scholarship (PGSD2-487305-2016, to E.M.A) and by a National Institutes of Health Award (1R01-GM121498, to W.R.H.).

