## Supplemental tables and figures for "Cross-feeding modulates the rate and mechanism of antibiotic resistance evolution in a model microbial community of *Escherichia coli* and *Salmonella enterica*"

**Supplementary tables and figures**

**Supplementary table 1. Mutation list.**

| **Antibiotic** | **Evolution condition** | **Replicate** | **Library** | **Species** | **Gene** | **Description** | **Mutation** | **Position** | **Freq** |
| --- | --- | --- | --- | --- | --- | --- | --- | --- | --- |
| Rifampicin | *E. coli* monoculture | 1 | BA010 | *E. coli* | *mdoH →* | glucosyltransferase | +T | 1,107,543:1 | 94.10% |
| Rifampicin | *E. coli* monoculture | 1 | BA010 | *E. coli* | *prc ←* | tail‑specific protease | Δ2 bp | 1,908,859 | 100% |
| Rifampicin | *E. coli* monoculture | 1 | BA010 | *E. coli* | *rpoB →* | DNA‑directed RNA polymerase subunit beta | A→C | 4,172,886 | 93.90% |
| Rifampicin | *E. coli* monoculture | 2 | BA017 | *E. coli* | *rpoB →* | DNA‑directed RNA polymerase subunit beta | C→T | 4,172,863 | 100% |
| Rifampicin | *E. coli* monoculture | 3 | BA030 | *E. coli* | *mdoG →* | glucan biosynthesis protein G mdoG → | +TT | 1,106,042:1 | 100% |
| Rifampicin | *E. coli* monoculture | 3 | BA030 | *E. coli* | *rpoB →* | DNA‑directed RNA polymerase subunit beta | G→A | 4,172,781 | 100% |
| Rifampicin | *E. coli* monoculture | 4 | BA036 | *E. coli* | *glpA →* | anaerobic glycerol‑3‑phosphate dehydrogenase subunit A | +CTGCGCGGG | 2,346,218 | 100% |
| Rifampicin | *E. coli* monoculture | 4 | BA036 | *E. coli* | *mdoG →* | glucan biosynthesis protein G mdoG → | G→C | 1,105,310 | 64.80% |
| Rifampicin | *E. coli* monoculture | 4 | BA036 | *E. coli* | *prc ←* | tail‑specific protease | Δ11 bp | 1,907,811 | 100% |
| Rifampicin | *E. coli* monoculture | 5 | BA049 | *E. coli* | *mdoG →* | glucan biosynthesis protein G mdoG → | Δ8 bp | 1,105,573 | 100% |
| Rifampicin | *E. coli* monoculture | 5 | BA049 | *E. coli* | *rpoB →* | DNA‑directed RNA polymerase subunit beta | A→G | 4,172,719 | 100% |
| Rifampicin | *E. coli* monoculture | 6 | BA056 | *E. coli* | *fre →* | NAD(P)H‑flavin reductase | Δ13 bp | 4,019,988 | 86.50% |
| Rifampicin | *E. coli* monoculture | 6 | BA056 | *E. coli* | *mdoH →* | glucosyltransferase | Δ6 bp | 1,107,321 | 100% |
| Rifampicin | *S. enterica* monoculture | 1 | BA067 | *S. enterica* | *mdoH →* | gucans biosynthesis glucosyltransferase H | G→T | 1,239,272 | 100% |
| Rifampicin | *S. enterica* monoculture | 1 | BA067 | *S. enterica* | *rpoB →* | DNA‑directed RNA polymerase subunit beta | T→G | 4,367,622 | 100% |
| Rifampicin | *S. enterica* monoculture | 2 | BA079 | *S. enterica* | *ispD ←* | 2‑C‑methyl‑D‑erythritol 4‑phosphate cytidylyltransferase | C→T | 3,070,986 | 100% |
| Rifampicin | *S. enterica* monoculture | 2 | BA079 | *S. enterica* | *mdoH →* | gucans biosynthesis glucosyltransferase H | G→T | 1,239,272 | 100% |
| Rifampicin | *S. enterica* monoculture | 2 | BA079 | *S. enterica* | *rpoB →* | DNA‑directed RNA polymerase subunit beta | A→G | 4,367,454 | 100% |
| Rifampicin | *S. enterica* monoculture | 2 | BA079 | *S. enterica* | *STM4466 ←* | carbamate kinase | A→G | 4,708,815 | 100% |
| Rifampicin | *S. enterica* monoculture | 4 | BA098 | *S. enterica* | *mdoH →* | gucans biosynthesis glucosyltransferase H | G→T | 1,239,272 | 100% |
| Rifampicin | *S. enterica* monoculture | 4 | BA098 | *S. enterica* | *rpoB →* | DNA‑directed RNA polymerase subunit beta | C→T | 4,367,442 | 100% |
| Rifampicin | *S. enterica* monoculture | 5 | BA108 | *S. enterica* | *rpoB →* | DNA‑directed RNA polymerase subunit beta | C→T | 4,367,511 | 90.00% |
| Rifampicin | *S. enterica* monoculture | 5 | BA108 | *S. enterica* | *rpoB →* | DNA‑directed RNA polymerase subunit beta | T→G | 4,367,622 | 100% |
| Rifampicin | *S. enterica* monoculture | 5 | BA108 | *S. enterica* | *ramR ←* | regulatory protein | Δ4 bp | 638,200 | 86.10% |
| Rifampicin | *S. enterica* monoculture | 6 | BA115 | *S. enterica* | *mdoH →* | glucans biosynthesis glucosyltransferase H | G→T | 1,239,272 | 100% |
| Rifampicin | *S. enterica* monoculture | 6 | BA115 | *S. enterica* | *rpoB →* | DNA‑directed RNA polymerase subunit beta | C→T | 4,367,504 | 100% |
| Rifampicin | Co-culture | 1 | BA125 | *E. coli* | *pnp ←* | polyribonucleotide nucleotidyltransferase | +G | 3,303,449:1 | 90.10% |
| Rifampicin | Co-culture | 1 | BA125 | *E. coli* | *rpoB →* | DNA‑directed RNA polymerase subunit betat | T→A | 4,172,887 | 100% |
| Rifampicin | Co-culture | 2 | BA139 | *E. coli* | *rfaQ ←* | LPS core heptosyltransferase RfaQ | Δ1 bp | 3,800,489 | 100% |
| Rifampicin | Co-culture | 2 | BA139 | *E. coli* | *rpoB →* | DNA‑directed RNA polymerase subunit beta | C→A | 4,172,893 | 100% |
| Rifampicin | Co-culture | 2 | BA139 | *S. enterica* | *rpoB →* | DNA‑directed RNA polymerase subunit beta | A→G | 4,366,350 | 100% |
| Rifampicin | Co-culture | 3 | BA145 | *E. coli* | *prc ←* | tail‑specific protease | Δ2 bp | 1,908,859 | 100% |
| Rifampicin | Co-culture | 3 | BA145 | *E. coli* | *prs ←* | ribose‑phosphate pyrophosphokinase | A→T | 1,256,880 | 100% |
| Rifampicin | Co-culture | 3 | BA145 | *E. coli* | *rpoB →* | DNA‑directed RNA polymerase subunit beta | C→A | 4,172,893 | 100% |
| Rifampicin | Co-culture | 4 | BA155 | *E. coli* | *prc ←* | tail‑specific protease | Δ10 bp | 1,907,831 | 64.40% |
| Rifampicin | Co-culture | 5 | BA165 | *E. coli* | *rplK →* | 50S ribosomal protein L11 | C→T | 4,168,451 | 100% |
| Rifampicin | Co-culture | 5 | BA165 | *E. coli* | *rpoB →* | DNA‑directed RNA polymerase subunit beta | C→A | 4,172,893 | 92.80% |
| Rifampicin | Co-culture | 5 | BA165 | *E. coli* | *BW25113_RS13710* ←  / ← *BW25113_RS13715* | CP4‑57 defective prophage, DUF4297/DUF1837  polymorphic toxin family protein/hypothetical protein | +GCACTATG | 2,758,778 | 87.60% |
| Rifampicin | Co-culture | 6 | BA176 | *E. coli* | *prc ←* | tail‑specific protease | Δ11 bp | 1,908,656 | 100% |
| Rifampicin | Co-culture | 6 | BA176 | *S. enterica* | *rpoB →* | DNA‑directed RNA polymerase subunit beta | C→A | 4,367,483 | 100% |
| Ampicillin | *E. coli* monoculture | 1 | rMM010 | *E. coli* | *acrB ←* | multidrug efflux RND transporter permease subunit | A→C | 479,480 | 100% |
| Ampicillin | *E. coli* monoculture | 1 | rMM010 | *E. coli* | *rne ←* | ribonuclease E | repeat_region (+) +5 bp :: Δ1 bp | 1,138,341 | 100% |
| Ampicillin | *E. coli* monoculture | 2 | rMM020 | *E. coli* | *envZ ←* | two‑component sensor histidine kinase | C→G | 3,528,288 | 100% |
| Ampicillin | *E. coli* monoculture | 4 | rMM039 | *E. coli* | *mdoH →* | glucosyltransferase | Δ1 bp | 1,107,469 | 100% |
| Ampicillin | *E. coli* monoculture | 5 | rMM049 | *E. coli* | *ilvN ←* | acetolactate synthase isozyme 1 small subunit | Δ5 bp | 3,844,420 | 72.20% |
| Ampicillin | *E. coli* monoculture | 6 | rMM060 | *E. coli* | *eda ←* | 2‑keto‑3‑deoxy‑L‑rhamnonate aldolase | Δ47 bp | 2,351,941 | 67.60% |
| Ampicillin | *E. coli* monoculture | 6 | rMM060 | *E. coli* | *ompF ←* | outer membrane protein F | Δ2 bp | 982,235 | 100% |
| Ampicillin | *E. coli* monoculture | 6 | rMM060 | *E. coli* | *prlF →* | antitoxin PrlF | repeat_region (–) +4 bp :: Δ3 bp | 3,270,368 | 59.20% |
| Ampicillin | *E. coli* monoculture | 6 | rMM060 | *E. coli* | *rne ←* | ribonuclease E | repeat_region (+) +5 bp :: Δ1 bp | 1,138,341 | 100% |
| Ampicillin | *S. enterica* monoculture | 1 | rMM067 | *S. enterica* | *ompF/IS10* | outer membrane protein F/ repeat region | IS element insertion | 1090025 =  = 1090033 | 85% |
| Ampicillin | *S. enterica* monoculture | 2 | rMM078 | *S. enterica* | *ompF/IS10* | outer membrane protein F/ repeat region | IS element insertion | 1090025 =  = 1090033 | 95% |
| Ampicillin | *S. enterica* monoculture | 2 | rMM078 | *S. enterica* | *ramR ←* | regulatory protein | coding (511‑554/582 nt) | 638,174 | 100% |
| Ampicillin | *S. enterica* monoculture | 2 | rMM078 | *S. enterica* | *STM2273/ IS10* | dehydratase/ repeat region | IS element insertion | 2377331 =  = 2377339 | 98% |
| Ampicillin | *S. enterica* monoculture | 3 | rMM090 | *S. enterica* | *acrB ←* | RND family acridine efflux pump | W634R (TGG→CGG) | 530,497 | 100% |
| Ampicillin | *S. enterica* monoculture | 3 | rMM090 | *S. enterica* | *ompR ←* | osmolarity response regulator OmpR | R210L (CGT→CTT) | 3,659,697 | 100% |
| Ampicillin | *S. enterica* monoculture | 3 | rMM090 | *S. enterica* | *ramR ←* | regulatory protein | Q19* (CAG→TAG) | 638,673 | 100% |
| Ampicillin | *S. enterica* monoculture | 3 | rMM090 | *S. enterica* | *STM2273/ IS10* | dehydratase/ repeat region | IS element insertion | 2377331 =  = 2377339 | 98% |
| Ampicillin | *S. enterica* monoculture | 3 | rMM090 | *S. enterica* | *yoaE* | inner membrane protein | Intragenic inversion | = 1926896 | 100% |
| Ampicillin | *S. enterica* monoculture | 4 | rMM098 | *S. enterica* | *STM2273/ IS10* | dehydratase/ repeat region | IS element insertion | 2377331 =  = 2377339 | 98% |
| Ampicillin | *S. enterica* monoculture | 5 | rMM108 | *S. enterica* | *ompF/IS10* | outer membrane protein F/ repeat region | IS element insertion | 1090025 =  = 1090033 | 73% |
| Ampicillin | *S. enterica* monoculture | 5 | rMM108 | *S. enterica* | *STM2273/ IS10* | dehydratase/ repeat region | IS element insertion | 2377331 =  = 2377339 | 98% |
| Ampicillin | *S. enterica* monoculture | 5 | rMM108 | *S. enterica* | *yoaE* | inner membrane protein | Intragenic inversion | = 1926896 | 84% |
| Ampicillin | *S. enterica* monoculture | 6 | rMM119 | *S. enterica* | *acrB ←* | RND family acridine efflux pump | F615S (TTC→TCC) | 530,553 | 100% |
| Ampicillin | *S. enterica* monoculture | 6 | rMM119 | *S. enterica* | *ompR ←* | osmolarity response regulator OmpR | R210L (CGT→CTT) | 3,659,697 | 100% |
| Ampicillin | Co-culture | 1 | rMM127 | *S. enterica* | *ahpF →* | alkyl hydroperoxide reductase subunit F | G→A | 672,700 | 94.60% |
| Ampicillin | Co-culture | 1 | rMM127 | *S. enterica* | *amn ←* | AMP nucleosidase | T→C | 2,092,111 | 100% |
| Ampicillin | Co-culture | 1 | rMM127 | *S. enterica* | *dnaQ →* | DNA polymerase III subunit epsilon | T→A | 303,499 | 100% |
| Ampicillin | Co-culture | 1 | rMM127 | *S. enterica* | *envZ ←* | osmolarity sensor protein EnvZ | A→G | 3,659,359 | 100% |
| Ampicillin | Co-culture | 1 | rMM127 | *S. enterica* | *ftsI →* | peptidoglycan synthase FtsI | +A | 143,219:1 | 100% |
| Ampicillin | Co-culture | 1 | rMM127 | *S. enterica* | *ftsZ →* | cell division protein FtsZ | C→T | 155,877 | 100% |
| Ampicillin | Co-culture | 1 | rMM127 | *S. enterica* | *gldA ← / → STM3531* | glycerol dehydrogenase/  dihydroxyacid dehydratase | G→A | 3,694,118 | 94.40% |
| Ampicillin | Co-culture | 1 | rMM127 | *S. enterica* | *metL →* | bifunctional aspartate kinase II/  homoserine dehydrogenase II | G→A | 4,312,839 | 100% |
| Ampicillin | Co-culture | 1 | rMM127 | *S. enterica* | *rtn →* | lambda/N4 phages resistance membrane protein | T→C | 2,315,609 | 92.40% |
| Ampicillin | Co-culture | 1 | rMM127 | *S. enterica* | *sppA ←* | protease 4 | G→A | 1,373,495 | 54.20% |
| Ampicillin | Co-culture | 1 | rMM127 | *S. enterica* | *STM0019 →* | hydroxymethyltransferase | A→G | 20,208 | 100% |
| Ampicillin | Co-culture | 1 | rMM127 | *S. enterica* | *STM0566 →* | inner membrane protein | G→A | 622,193 | 100% |
| Ampicillin | Co-culture | 1 | rMM127 | *S. enterica* | *STM1552 → / ← STM05155* | cytoplasmic protein/hypothetical protein | A→G | 1,629,730 | 100% |
| Ampicillin | Co-culture | 1 | rMM127 | *S. enterica* | *STM2179 ←* | sugar transporter | T→C | 2,275,700 | 93.00% |
| Ampicillin | Co-culture | 1 | rMM127 | *S. enterica* | *STM2700 ←* | phage tail fiber‑like protein | T→C | 2,850,036 | 94.00% |
| Ampicillin | Co-culture | 1 | rMM127 | *S. enterica* | *STM2739 →* | phage tail‑like protein | C→A | 2,877,206 | 100% |
| Ampicillin | Co-culture | 1 | rMM127 | *S. enterica* | *STM2756 ←* | sugar phosphate aminotransferase | C→T | 2,894,787 | 100% |
| Ampicillin | Co-culture | 1 | rMM127 | *S. enterica* | *STM3052 ←* | outer membrane protein | T→C | 3,211,576 | 94.00% |
| Ampicillin | Co-culture | 1 | rMM127 | *S. enterica* | *STM3631 ←* | xanthine permease | T→C | 3,817,700 | 94.60% |
| Ampicillin | Co-culture | 1 | rMM127 | *S. enterica* | *STM3653 ← / ← glyS* | acetyltransferase/glycine‑‑tRNA ligase subunit beta | A→G | 3,839,640 | 100% |
| Ampicillin | Co-culture | 1 | rMM127 | *S. enterica* | *STM4419 →* | sugar transporter | C→T | 4,662,084 | 100% |
| Ampicillin | Co-culture | 1 | rMM127 | *S. enterica* | *xylA ← / → xylR* | xylose isomerase/xylose operon regulatory protein | T→C | 3,848,052 | 100% |
| Ampicillin | Co-culture | 1 | rMM127 | *S. enterica* | *yeaQ →* | inner membrane protein | T→C | 1,353,286 | 54.60% |
| Ampicillin | Co-culture | 1 | rMM127 | *S. enterica* | *yhiP →* | dipeptide/tripeptide permease B | C→T | 3,762,685 | 52.60% |
| Ampicillin | Co-culture | 2 | rMM137 | *E. coli* | *proQ ←* | RNA chaperone ProQ | Δ5 bp | 1,909,389 | 19.70% |
| Ampicillin | Co-culture | 2 | rMM137 | *S. enterica* | *metL →* | bifunctional aspartate kinase II/  homoserine dehydrogenase II | Δ4 bp | 4,311,847 | 59.20% |
| Ampicillin | Co-culture | 3 | rMM146 | *S. enterica* | *ompF ←* | outer membrane protein F | Δ116 bp | 1,090,110 | 64.50% |
| Ampicillin | Co-culture | 4 | rMM158 | *E. coli* | *proQ ←* | RNA chaperone ProQ | C→A | 1,909,737 | 86.50% |
| Ampicillin | Co-culture | 4 | rMM158 | *S. enterica* | *ftsI →* | peptidoglycan synthase FtsI | G→T | 143,325 | 75.30% |
| Ampicillin | Co-culture | 5 | rMM167 | *S. enterica* | *yoaE* | inner membrane protein | Intragenic inversion | = 1926896 | 100% |

Supplementary table 2. Gene functions

| Gene | Mutated in | Description | Function |
| --- | --- | --- | --- |
| *fre →* | *E. coli*: BA056 | NAD(P)H‑flavin reductase | May be involved in iron homeostasis and oxidative stress response (UniProt) |
| *rpoB →* | *E. coli*: BA010, BA017, BA030, BA049, BA125, BA139, BA145, BA165  *S. enterica*: BA067, BA079, BA098, BA108, BA115, BA139, BA176 | DNA‑directed RNA polymerase subunit beta | RNA polymerase beta subunit; commonly mutated in rifampicin-resistant lines (1) |
| *prc ←* | *E. coli*: BA010, BA036, BA145, BA155, BA176 | tail‑specific protease | Cleaves precursor to form a functional PBP3; involved in thermal and osmotic stress response (2); mutations lead to increased antibiotic susceptibility (3) |
| *mdoH →* | *E. coli*: BA010, BA056, rMM039  *S. enterica*: BA067, BA079, BA098, BA115 | glucosyltransferase | Also called *mdoH/opgH*; involved in regulating cell wall osmolarity and likely modulates cellular penetration of rifampicin (4) |
| *mdoG →* | *E. coli*: BA030, BA036, BA049 | glucan biosynthesis protein G MdoG → | Required for the synthesis of osmoregulated periplasmic glucans; may function in biofilm formation (5) |
| *glpA →* | *E. coli*: BA036 | anaerobic glycerol‑3‑phosphate dehydrogenase subunit A | Involved in utilization of glycerol as a carbon source; deletions lead to decreased persister formation (6) |
| *pnp ←* | *E. coli*: BA125 | polyribonucleotide nucleotidyltransferase | Involved in mRNA degradation and tRNA processing; contributes to rRNA quality control during steady-state growth (7) |
| *rfaQ ←* | *E. coli*: BA139 | LPS core heptosyltransferase RfaQ | Also called *waaQ*; involved in LPS biosynthesis (UniProt) |
| *prs ←* | *E. coli*: BA145 | ribose‑phosphate pyrophosphokinase | Involved in central metabolism (UniProt) |
| BW25113_RS13710 ←  / ← BW25113_RS13715 | *E. coli*: BA165 | CP4‑57 defective prophage, DUF4297/DUF1837 polymorphic toxin family protein/hypothetical protein |  |
| *rplK →* | *E. coli*: BA165 | 50S ribosomal protein L11 | Regulator of the stringent response and signals increased ppGpp production, which increases antibiotic tolerance (8) |
| *rne ←* | *E. coli*: rMM010, rMM060 | ribonuclease E | Small regulatory RNA that functions in SOS initiation (9) |
| *acrB ←* | *E. coli*: rMM010  *S. enterica*: rMM090, rMM119 | multidrug efflux RND transporter permease subunit | Efflux transporter protein component of the TolC-AcrAB multidrug efflux pump (10) |
| *envZ ←* | *E. coli*: rMM020  *S. enterica*: rMM127 | two‑component sensor histidine kinase | Sensor kinase in two-component signalling control of *ompF/ompC* expression regulation (10) |
| *ilvN ←* | *E. coli*: rMM049 | acetolactate synthase isozyme 1 small subunit | Catalyzes the first step in valine biosynthesis and the second step in isoleucine biosynthesis (UniProt) |
| *eda ←* | *E. coli*: rMM060 | 2‑keto‑3‑deoxy‑L‑rhamnonate aldolase | Involved in glucose degradation through the Entner-Doudoroff pathway (UniProt) |
| *prlF →* | *E. coli*: rMM060 | antitoxin PrlF | Antitoxin component of an mRNA degradation toxin system (11) |
| *ompF ←* | *E. coli*: rMM060  *S. enterica*: rMM146 | outer membrane protein F | Classic trimeric porin commonly lost in beta-lactam resistant strains (12) |
| *proQ ←* | *E. coli*: rMM137, rMM158 | RNA chaperone ProQ | Small regulatory RNA that controls efflux pump expression (13) |
| *ispD ←* | *S. enterica*: BA079 | 2‑C‑methyl‑D‑erythritol 4‑phosphate cytidylyltransferase | Functions in isoprene biosynthesis (UniProt) |
| *STM4466 ←* | *S. enterica*: BA079 | carbamate kinase | Functions in the arginine deaminase pathway (UniProt) |
| *ramR ←* | *S. enterica*: BA108, rMM078, rMM090 | regulatory protein | Negative repressor of *ramA*, which positively regulates *acrAB* expression- knockouts result in constitutive *acrAB* expression (14) |
| *ompF/IS10* | *S. enterica*: rMM067, rMM078, rMM108 | outer membrane protein F/ repeat region | Classic trimeric porin commonly lost in beta-lactam resistant strains (12) |
| *STM2273/ IS10* | *S. enterica*: rMM078, rMM090, rMM098, rMM108 | dehydratase/ repeat region | - |
| *yoaE* | *S. enterica*: rMM090, rMM108, rMM167 | inner membrane protein | Integral membrane protein, putative transporter (UniProt) |
| *ompR ←* | *S. enterica*: rMM090, rMM119 | osmolarity response regulator OmpR | Response regulator in two-component signalling of *ompF/ompC* expression (10) |
| *ahpF →* | *S. enterica*: rMM127 | alkyl hydroperoxide reductase subunit F | Functions in protecting cells from hydrogen peroxide toxicity (15) |
| *amn ←* | *S. enterica*: rMM127 | AMP nucleosidase | Loss of function mutations allow greater cold tolerance in *E. coli* (16) |
| *dnaQ →* | *S. enterica*: rMM127 | DNA polymerase III subunit epsilon | Encodes proofreading DNA polymerase III; mutations lead to high mutation rates and are often observed under antibiotic selection (17) |
| *ftsZ →* | *S. enterica*: rMM127 | cell division protein FtsZ | Essential component of cell division (forms septum Z-ring); target of antimicrobial development (18) |
| *gldA ← / → STM3531* | *S. enterica*: rMM127 | glycerol dehydrogenase/dihydroxyacid dehydratase | - |
| *rtn →* | *S. enterica*: rMM127 | lambda/N4 phages resistance membrane protein | - |
| *sppA ←* | *S. enterica*: rMM127 | protease 4 | Signal peptide peptidase (UniProt) |
| *STM0019 →* | *S. enterica*: rMM127 | hydroxymethyltransferase | - |
| *STM0566 →* | *S. enterica*: rMM127 | inner membrane protein | - |
| *STM1552 → / ← STM05155* | *S. enterica*: rMM127 | cytoplasmic protein/hypothetical protein | - |
| *STM2179 ←* | *S. enterica*: rMM127 | sugar transporter | 4-hydroxybenzoate transporter (UniProt) |
| *STM2700 ←* | *S. enterica*: rMM127 | phage tail fiber‑like protein | - |
| *STM2739 →* | *S. enterica*: rMM127 | phage tail‑like protein | Putative integrase (UniProt) |
| *STM2756 ←* | *S. enterica*: rMM127 | sugar phosphate aminotransferase | - |
| *STM3052 ←* | *S. enterica*: rMM127 | outer membrane protein | May be involved in phenol degradation (UniProt) |
| *STM3631 ←* | *S. enterica*: rMM127 | xanthine permease | - |
| *STM3653 ← / ← glyS* | *S. enterica*: rMM127 | acetyltransferase/glycine‑‑tRNA ligase subunit beta | - |
| *STM4419 →* | *S. enterica*: rMM127 | sugar transporter | Carbohydrate/proton symporter (UniProt) |
| *xylA ← / → xylR* | *S. enterica*: rMM127 | xylose isomerase/xylose operon regulatory protein | - |
| *yeaQ →* | *S. enterica*: rMM127 | inner membrane protein | Putative integral membrane protein (UniProt) |
| *yhiP →* | *S. enterica*: rMM127 | dipeptide/tripeptide permease B | Putative transporter (UniProt) |
| *metL →* | *S. enterica*: rMM127, rMM137 | bifunctional aspartate kinase II/homoserine dehydrogenase II | Catalyzes the first step in lysine/homoserine biosynthesis, the last step in homoserine biosynthesis, and indirectly functions in methionine and threonine biosynthesis (UniProt) |
| *ftsI →* | *S. enterica*: rMM127, rMM158 | peptidoglycan synthase FtsI | Penicillin-binding protein 3 (PBP3); mutations confer β-lactam resistance (19) |

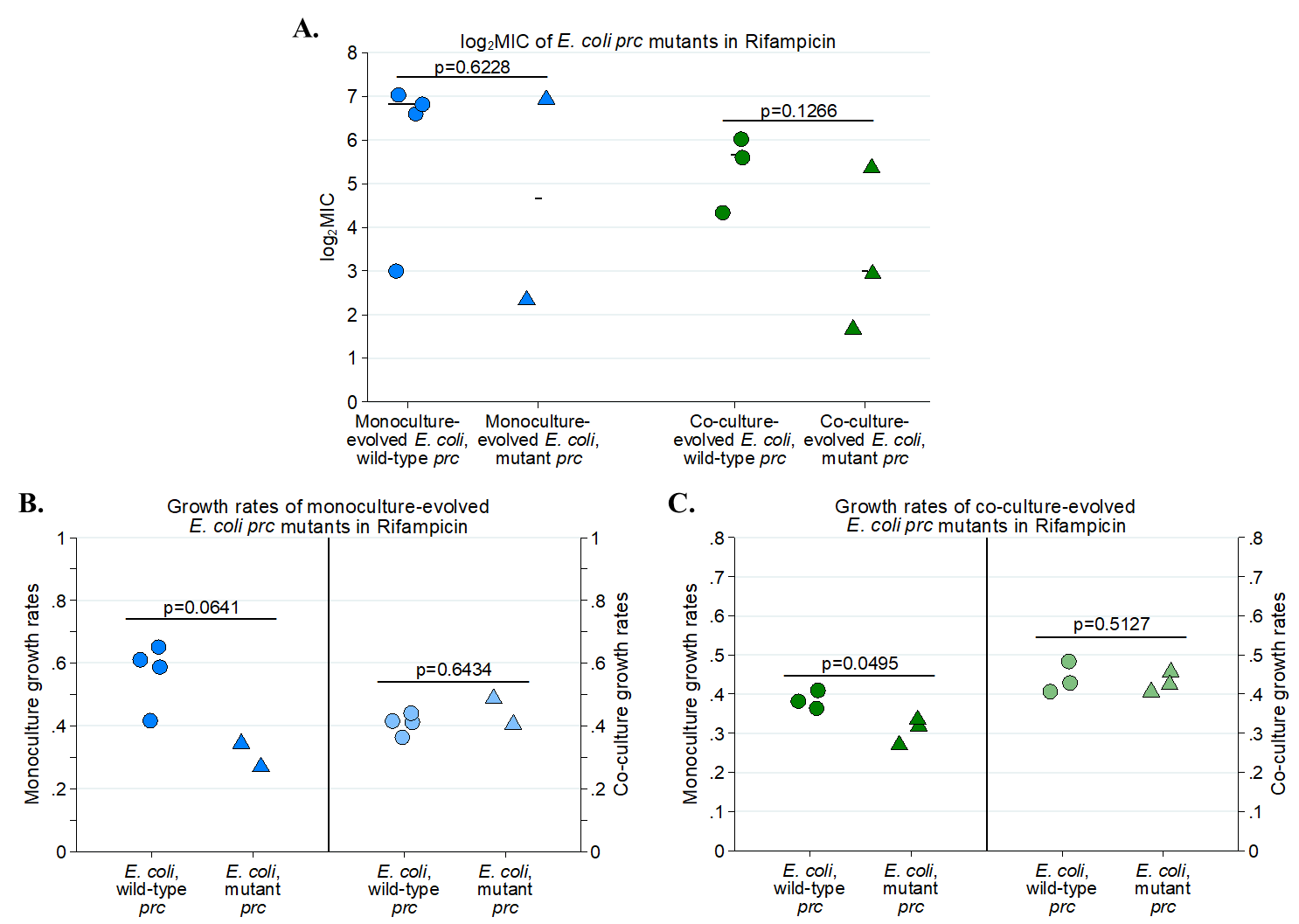

**Supplementary figure 1. A.** MICs of monoculture- and co-culture- evolved *E. coli* isolates containing wild-type or mutant *prc*. **B.** Monoculture and coculture growth rates of monoculture-evolved *E. coli* isolates containing wild-type or mutant *prc*. **C.** Monoculture and coculture growth rates of co-culture-evolved *E. coli* isolates containing wild-type or mutant *prc*. All p-values based on Mann-Whitney U tests. Points represent the average growth rate or MIC values of three isolates from the same population.

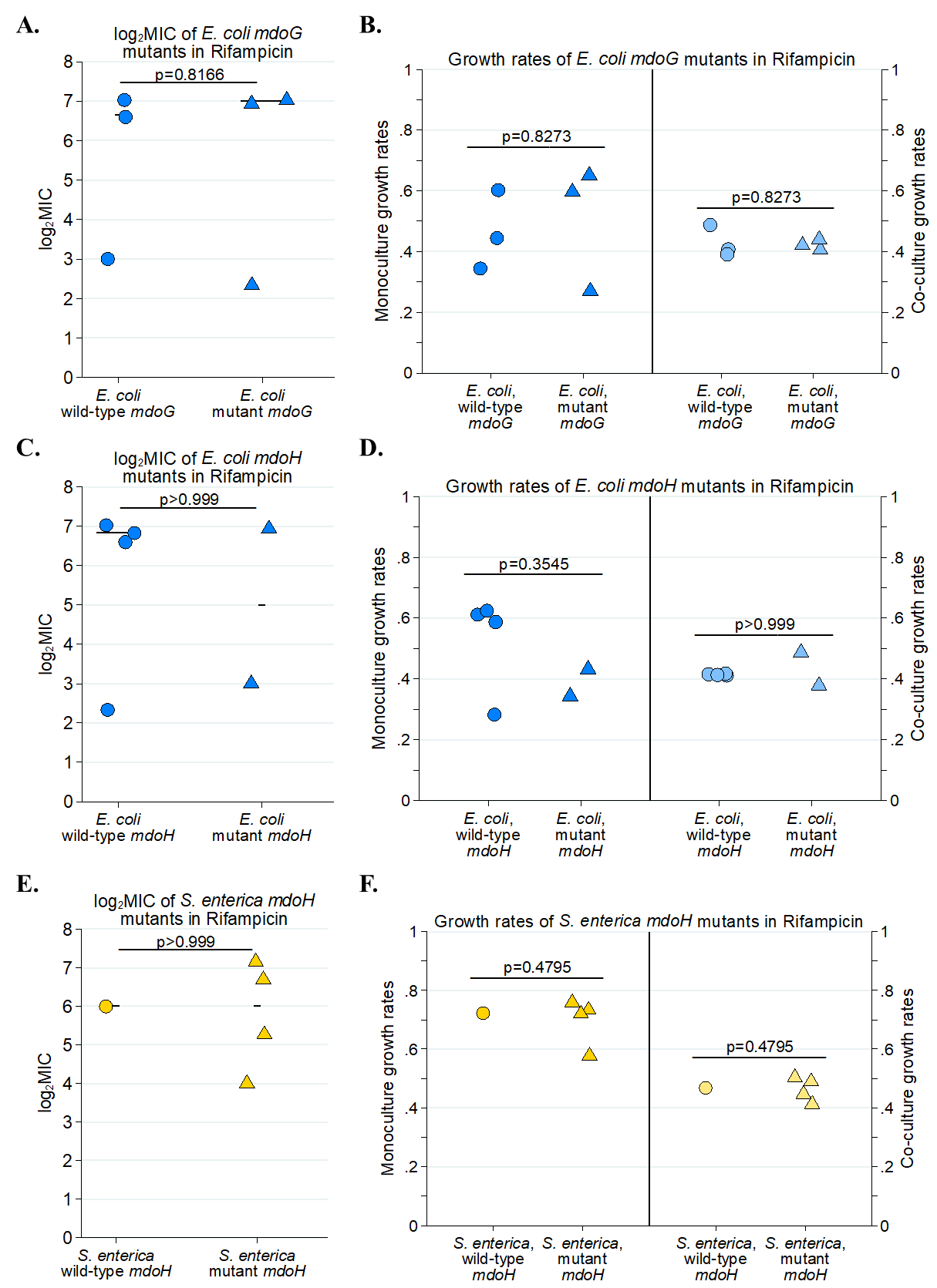

**Supplementary figure 2**. Impact of *mdoG* and *mdoH* mutations on MICs and growth rates in rifampicin-resistant evolved *E. coli* and *S. enterica*. **A.** MIC of *mdoG* wild-type vs. mutant *E. coli* isolates, averaged by population (three isolates per population were used). **B.** Monoculture and co-culture growth rates of *mdoG* wild-type vs. mutant *E. coli* isolates. **C.** MIC of *mdoH* wild-type vs. mutant *E. coli* isolates. **D.** Monoculture and co-culture growth rates of *mdoH* wild-type vs. mutant *E. coli* isolates. E**.** MIC of *mdoH* wild-type vs. mutant *S. enterica* isolates. **F.** Monoculture and co-culture growth rates of *mdoH* wild-type vs. mutant *S. enterica* isolates.

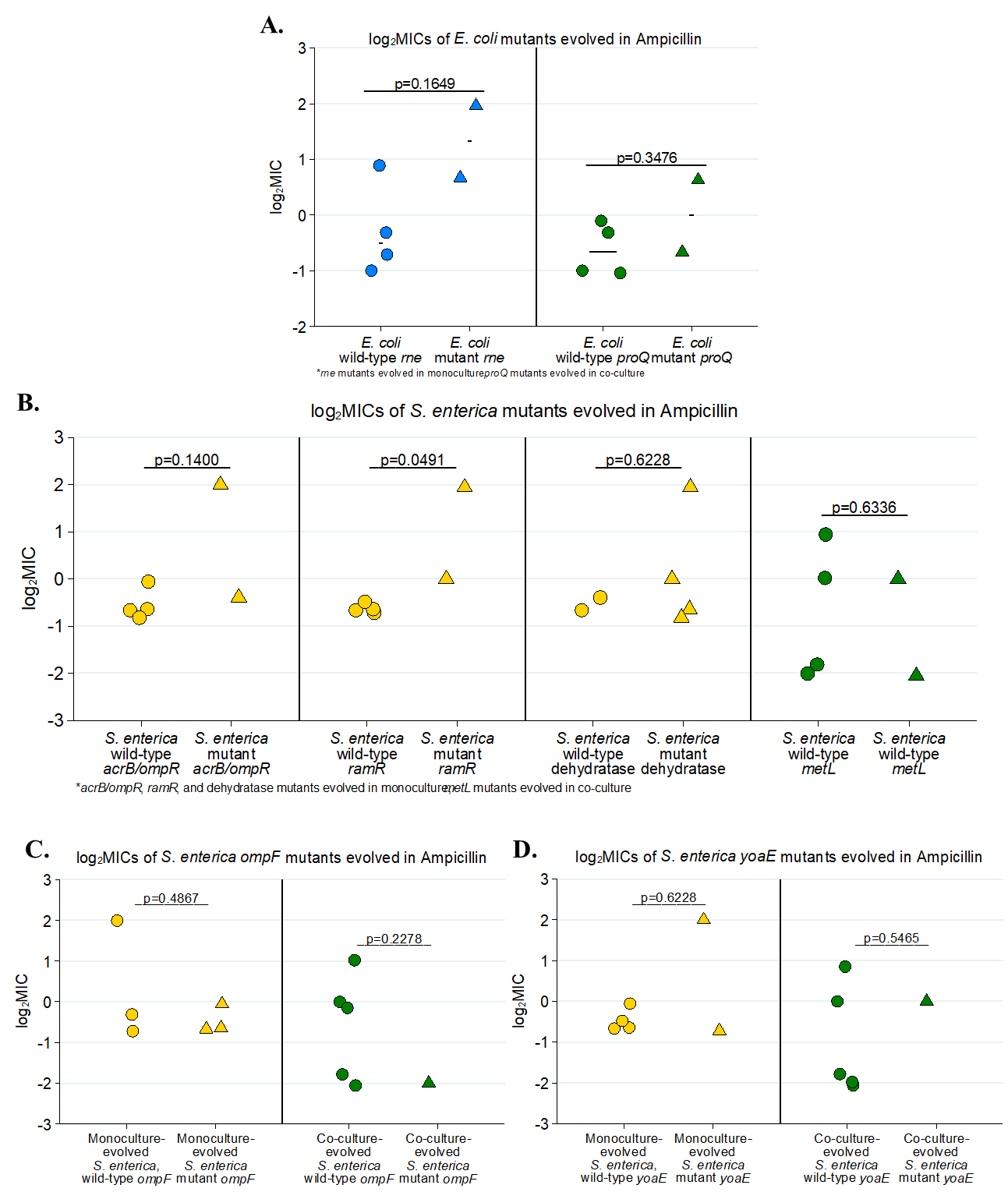

**Supplementary figure 3.** Effect of other mutations on MICs of ampicillin-evolved isolates. **A.** MICs of wild-type vs. mutant *E. coli* with mutations in *rne* (monoculture-evolved, in blue) and *proQ* (coculture-evolved, in green). **B.** MICs of wild-type vs. mutant *S. enterica* with mutations in *acrB/ompR*, *ramR,* an unnamed dehydratase (all evolved in monoculture only), and *metL* (evolved in co-culture only). **C.** MICs of wild-type vs. mutant *S. enterica* with mutations in *ompF* evolved in monoculture (gold) or co-culture (green). **D.** MICs of wild-type vs. mutant *S. enterica* with mutations in *yoaE* evolved in monoculture (gold) or co-culture (green).

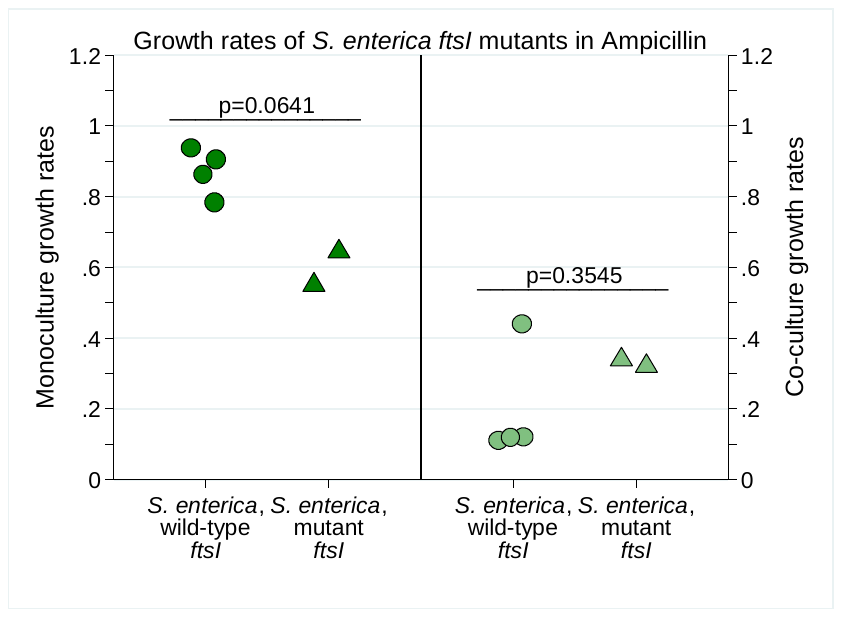

**Supplementary figure 4.** Monoculture and co-culture growth rates of *ftsI* mutant isolates in pH=4.7 growth medium. P= 0.0614 for monocultures, p= 0.3545 for co-cultures, Mann-Whitney U test.

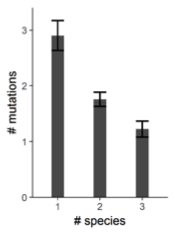

**Supplementary figure 5.** Average number of mutations that were observed in simulations with increasing numbers of species.
